## Supplementary figures and images for "High-Intensity Interval Training Remodels the Proteome and Acetylome of Human Skeletal Muscle"

### Figure 1 - figure supplement 1

#### CONSORT Flow Diagram

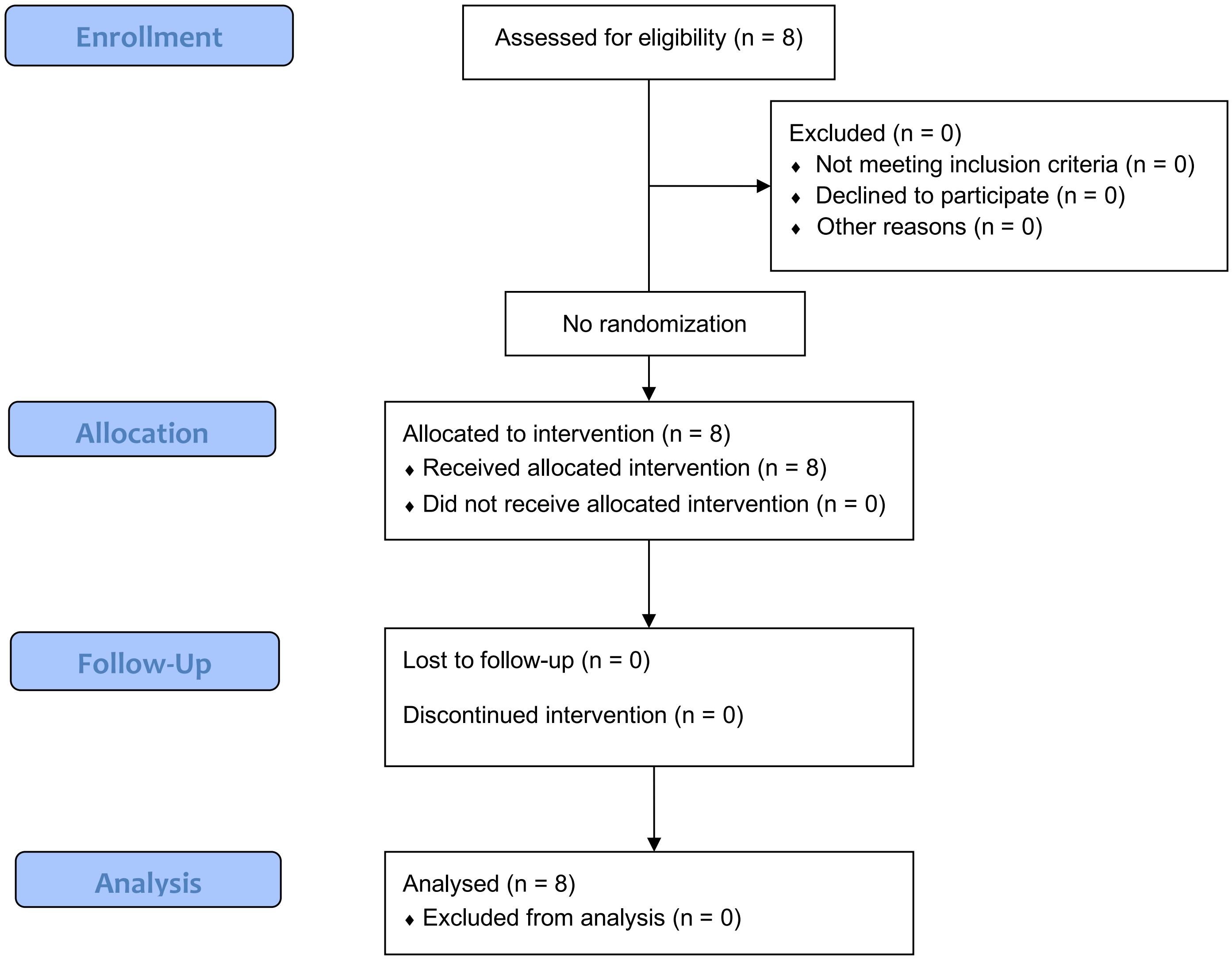

### Figure 1 - figure supplement 2

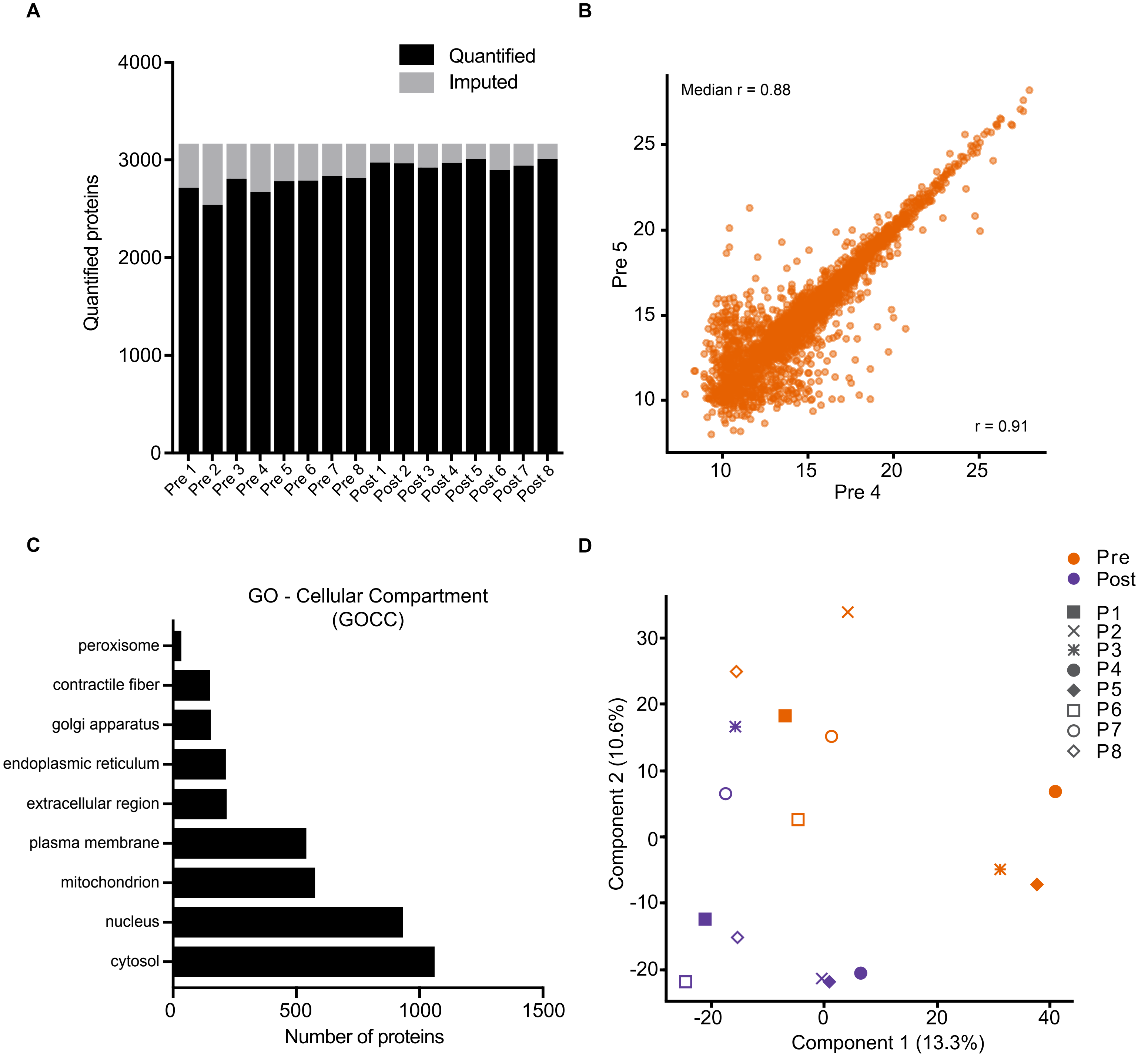

### Figure 2 - figure supplement 1

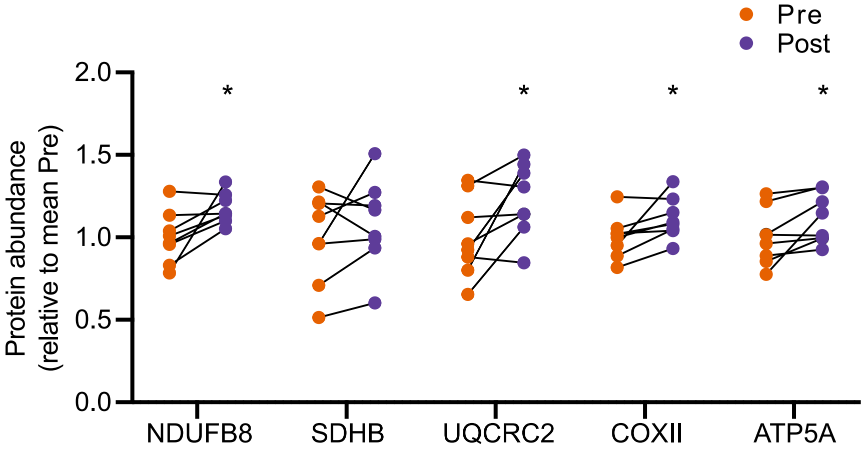

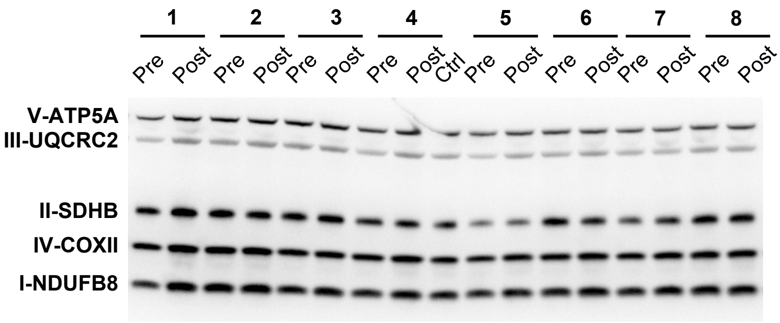

### Figure 3 - figure supplement 1

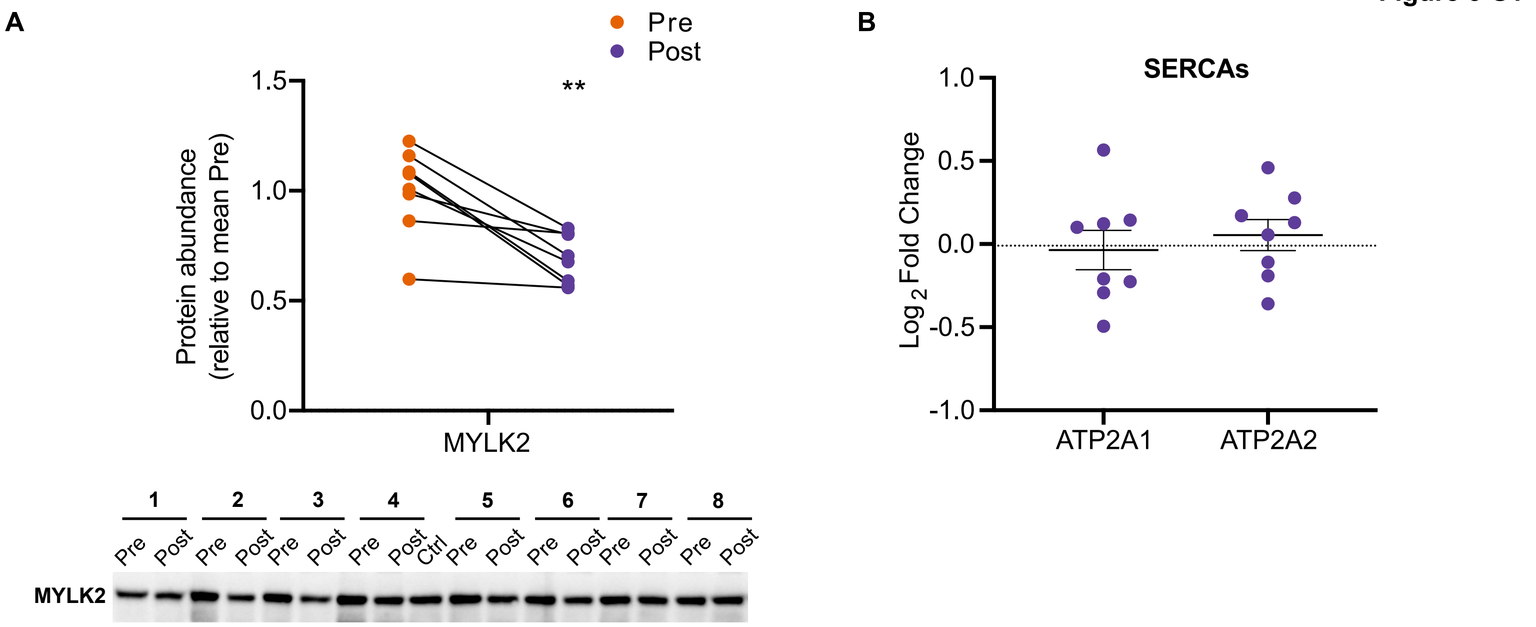

### Figure 4 - figure supplement 1

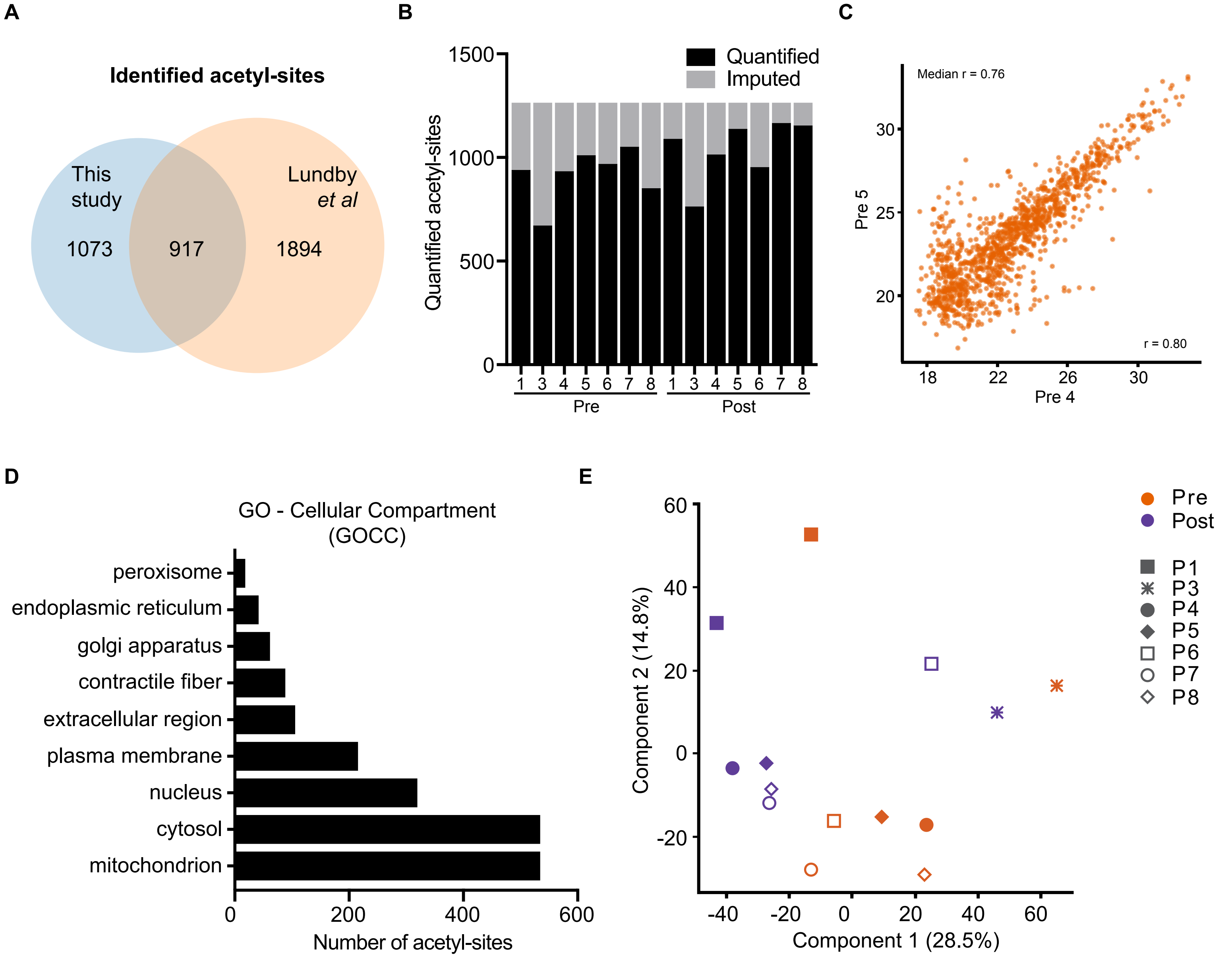

### Figure 5 - figure supplement 1

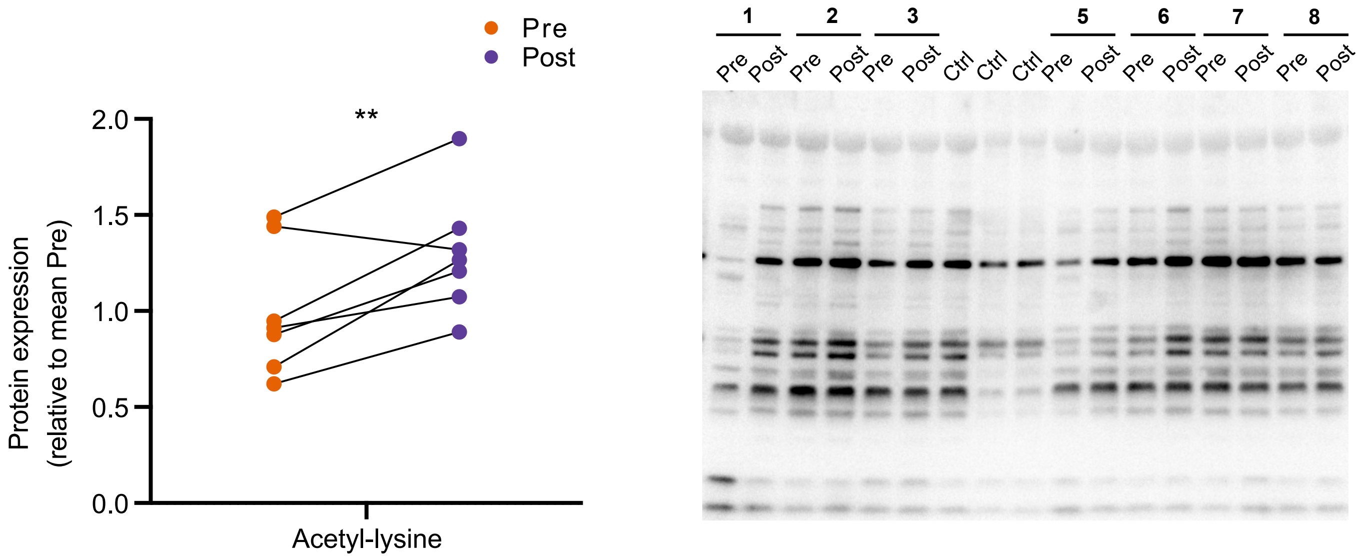

### Figure 5 - figure supplement 2

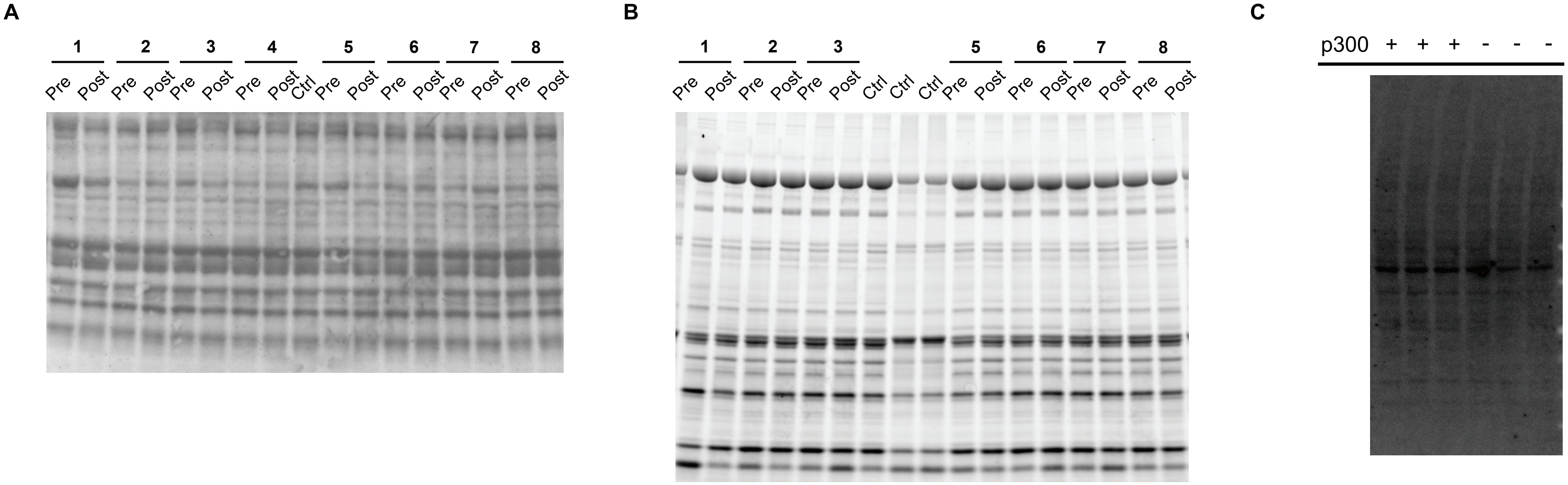
